# Comparative plant growth responses in a crop-weed mix – a case study of *Zea mays* (L.) and *Eleusine indica* (L.) Gaertn

**DOI:** 10.1101/2023.01.07.523107

**Authors:** Beckley Ikhajiagbe, Saheed Ibrahim Musa, Efuwa Famous Ekhator

## Abstract

The comparative plant growth responses of *Zea mays* and *Eleusine indica* were monitored in this studyunder an open field experiment.Both crop and weed were grown separately on the bowls and also in a crop-weed mix. Crop-weed mix were studied in different proportions. The experiment started first by acclimatizing the *E. indica* on the soil for a period of three days before sowing the maize. Forty (40) days after plant interactions, results showed significant decreases in plant heights and of both maize and *E. indica* depending on the ratio of both plants in the crop-weed mix.comparative to individual height. Similarly, significant decreases in leaf area was reported in *E. indica* (p=0.015) owing to the interaction. In the control, leaf area was 11.4cm^2^, but this decreased significantly to 5.94cm^2^ in the 2e3m mix. Similar decreases in maize leaf area have been reported with leaf area values being low (4.32cm^2)^ in the 5e3m mix compared to 19.9cm^2^ in the control (1m). Changes in prominent root length showed a significant decrease between the control *E. indica* and the crop-weed mix. Similar result was observed for maize plant. Although shoot – root ratio was 1.0 in the control weed (1e), a higher value (4.5) was observed in 1m1e followed by 3.5 in the 1m3e and 2m3e mix respectively. For maize plant however, the shoot-root ratio was 4.5 in the control (1m), but reduced to 1.5 in the 1e3m mix. The maize plant was also observed to release allelochemicals, which further indicate its competitive strength.

## Introduction

Invasion by exotic species is one of the most significant threats to biodiversity and food security globally (Alpert *et al*., 2000). Understanding the mechanisms by which invasive species such as weeds compete native species such as food crops is necessary to reduce the negative impacts of the invasive species, boost crop productivity and combat food insecurity. Also, understanding the which crop outcompete which weed and vice-versa may give information about plant physiology in weed stressed environments. The prevalence of severe food insecurity varies largely among different continents. Approximately 27.4% of Africa’s population was identified as severely food insecure in 2016, which is almost four times as high as any other region (FAO, 2017). Food insecurity is caused by a number of environmental factors such as the nutrient depletion as a results of invasive and native species competition, limit growth and security of crops. The agricultural interest of world is to obtain maximum crop yield, ensure food security and fulfill the food, feed and fiber requirements in order to meet with the ever increasing world population (Ikhajiagbe*et al*., 2019).

Crop is a plant that can be grown and harvested extensively for profit or subsistence.About 70% of the world’s agricultural land is given to crop grasses, and more than 50% of the world’s calories come from crops, especially the cereals (Conab, 2016). The agricultural interest of world aims at improving crop yield and fulfill the food, feed and fiber requirements in competition of fast increasing population. Most of these crops are found to produce edible grains which provides a rich source of carbohydrates for the germinating embryo (the germ). Proteins, oil and some vitamins are contained in the embryo (Ofor*et al*, 2009). Crops form a principal source of human food, such as wheat, rice, barely and maize being the widely consumed crops. These are often consumed for thousand years as staple food.

However, crop cultivation in most part of the world is currently facing problems ranging from management factors such as depth of sowing, row spacing, seed size and environmental stressors. Among all these, environmental stressors such as temperature, moisture, herbicides application, synthetic fertilizer usage presence of pests causing diseases and nutrient depletion has been considered common with tropical Africa regions. Soil nutrients can be depleted by a number of ways, weed inversion and competition is considered as the most frequent in tropical regions like Nigeria.

Weeds are a major environmental stressor and a major threat to the natural environment. It directly reduce crop yield and deteriorate its quality. One of the most common and invasive agricultural and environmental weed is *E. indica* commonly known as goose grass (Randall, 2012). It is considered as a serious weed in atleast42 countries including Nigeria (Holm *et al*., 2019; Waterhouse, 2011). The weed plant is tolerant of heavy disturbance and pollution and can grow along sewage lines, gutters very easily. Weed invasions create a change in the natural diversity and balance of ecological communities. These changes tend to threaten the survival rate of many plants since they compete with native plants for space, nutrients and sunlight. According to Ofor*et al*., 2009, weeds as pets significantly reduce productivity and survival of crops. According to Anchal*et al*., 2017; Azmi, 2000), rice crops faced about 70%yield reduction due to weed inversion and it is a higher yield loss than any other pest. Similarly, Oerke and Dehne (2004) have reported about 35% rice yield losses in DSR owing to weed competition worldwide. Baloch *et al*., 2012)observed that there was a low yield losses in sugarcane caused by high weed infestation.Weeds generally may out-compete native plants since they may not be affected by the pests or diseases that native crops would encounter in their natural habitats. Weeds can also reduce natural diversity and are capable of surviving and reproducing in disturbed environments and they often form the first species to colonize and dominate such environments in these conditions (Shad, 2019). Therefore devising a sustainable weed control method will help solve problems associated with crop productivity and food insecurity.

Several methods of weed control has been discussed by previous literatures. Chemical, mechanical, traditional and biological methods has been extensively studied (Shah *et al*., 2016). The biological methods has been suggested as the most striking, sustainable and eco-friendly. (Gibson *et al*., 2002). It involves strategies such as tillage, crop rotation and mixed cropping. Mixed cropping involves growing more than one crop at same field and time simultaneously which leads to competition and eventual death of the weak species (Kumar and Ijariya, 2013).The competition between crop–and-weed plants is governed by identifying which crops can compete with which weed, seed-weed rate and row spacing. The current research aimed at investigating investigate the morphological growth interactions of young maize plants and a common weed (*E. indica*). The study also considers the seed-weed rate and mix in the competitive setup. This research will enable researchers to explore the effectiveness of maize plant as competitive cultivar of weeds and if maize seed density affects invasive action of *E. indica*..

## Materials and Methods

### Experimental site

The study was conducted at the botanical garden of the department of Plant Biology and Biotechnology, Faculty of Life Sciences, University of Benin. The land was cleared off its overgrown weeds and a plot measuring 2.3m by 3m was marked out for the experiment.

### Soil sample preparation and seed collection

This begins by using a perforated 22 experimental bowls and each was filled up with soil obtained from biological garden at a depth of not more than 10cm. The maize seeds were obtainedfrom a harvested maize farm at Ovia North east local government area, Isihior, Benin City. *E. indica* was propagated using young tillers of 5 leaves each for the purpose of uniformity.

### Planting

The study was carried out under 11 treatments, with each one of them having a replicate. The treatments involved sowing of maize and *E. indica*separately as well as matching varying numbers of the crop and weed together.The young tillers of *E. indica* was first planted on the soil to acclimatize. After three days, the maize seeds were planted in the bowls according to the required treatments in order to monitor their interactions.After successful planting of both crop and weed, they were watered daily and various growth parameters were measured at weekly intervals.

### Measurement of experimental parameter

Plant based parameters such as plant height, number of leaves, number of additional tillers, leaf area, chlorophyll content, peduncle length, spike length, presence of chlorotic spot, prominent root length, number of roots, shoot to root length, dry weight of root and foliar colour were measured for each treatment following standard procedures. Survival percentage was calculated using the number of seed sown in each bowls and the number that survived at the end of the experiment. Presence of root acid of the both plant under the 11 treatments were determined using a blue litmus paper following FAO, 2007.

### Statistics

The mean and statistical error of data was calculated (Zar, 1974). Analysis of variance in complete by randomized design was done using the SPSS-16 statistical software, and means were separated by using the Least Significant Difference (Ogbeibu, 2005).

## Results and Discussion

Figure 1(a, b) presents information on plant height of maize, weed and their mixture for 40 days (Fig. 1a). For *E. indica*, plant height progressed from 5 cm on the 10^th^ day in the control experimental bowl with only the weed to 50 cm in same bowl 40 days later. When the weed was interacted with young maize plants at the rate of 1 weed to 5 maize plants (1e5m), plant height of *E. indica* progressed from 9.3cm to 30.2 cm at 40days. As well, plant height of maize without interaction with the weed increased from the 10^th^ day till 40th. However, within the same plant-weed mix, growth of maize plant increased from 2 cm on day 10 to 16.4cm on day 30, beyond which the plant died out (Fig. 1b). Similar plant die-outs in young maize plants were reported in 1m1e and 1m5e before the 40^th^ day after sowing. This showed that weed height is significantly inhibited with the interaction of maize plant and vice-versa showing competition. The death of maize plant before 40days showed invasive nature of *E*.*indica*with respect to maize due to prolonged period of interaction. This result is consistent with the work ofTunku and Ishaya (2012); Usoroh (2003) that keeping weeds with maize plant may lead to significant growth depression and possible death as a result of prolong interaction.

**Fig. 1(a):**
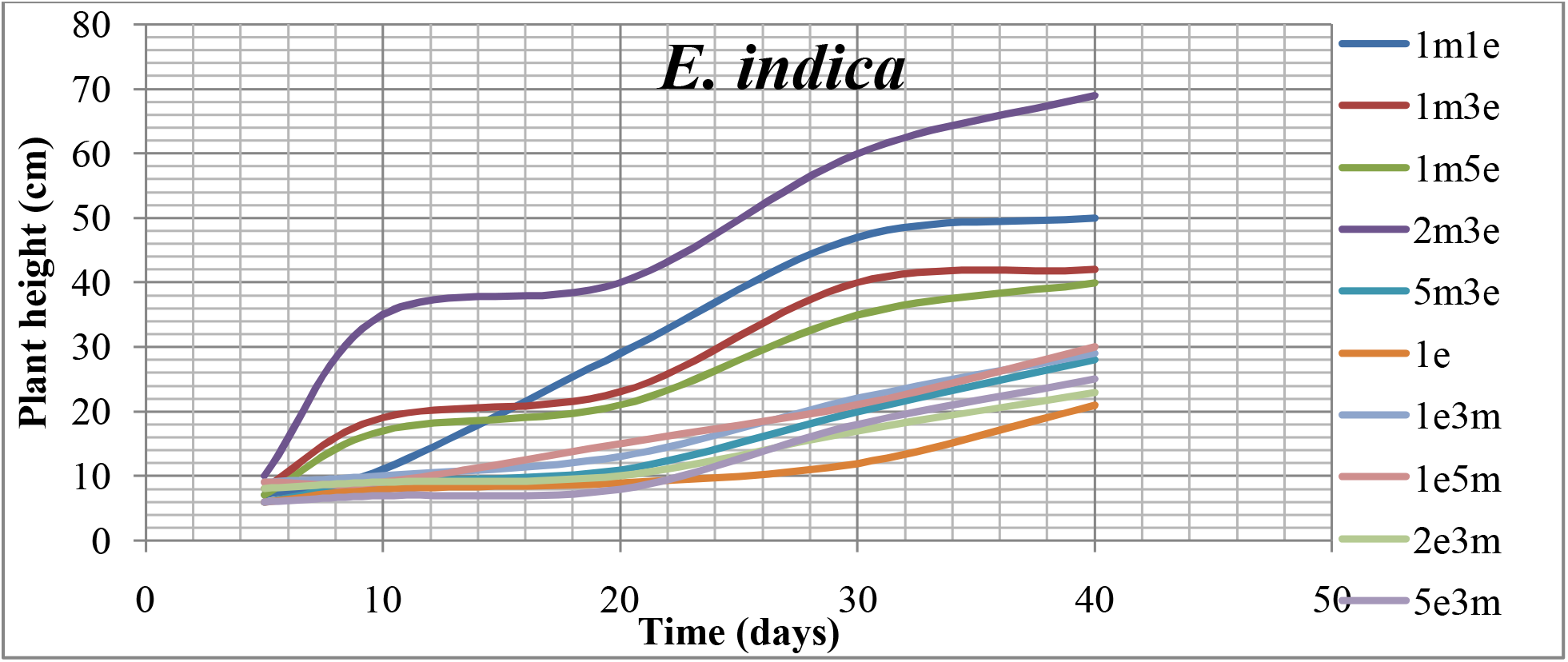
Effects of interaction between *Z. mays* and *E. indica* on plant height of *E. indica* (The plant interactions are represented on right hand side of graph, indicating number of *Eleusine* plants (e) in interaction with maize (m) plants)

**Fig. 1(b):**
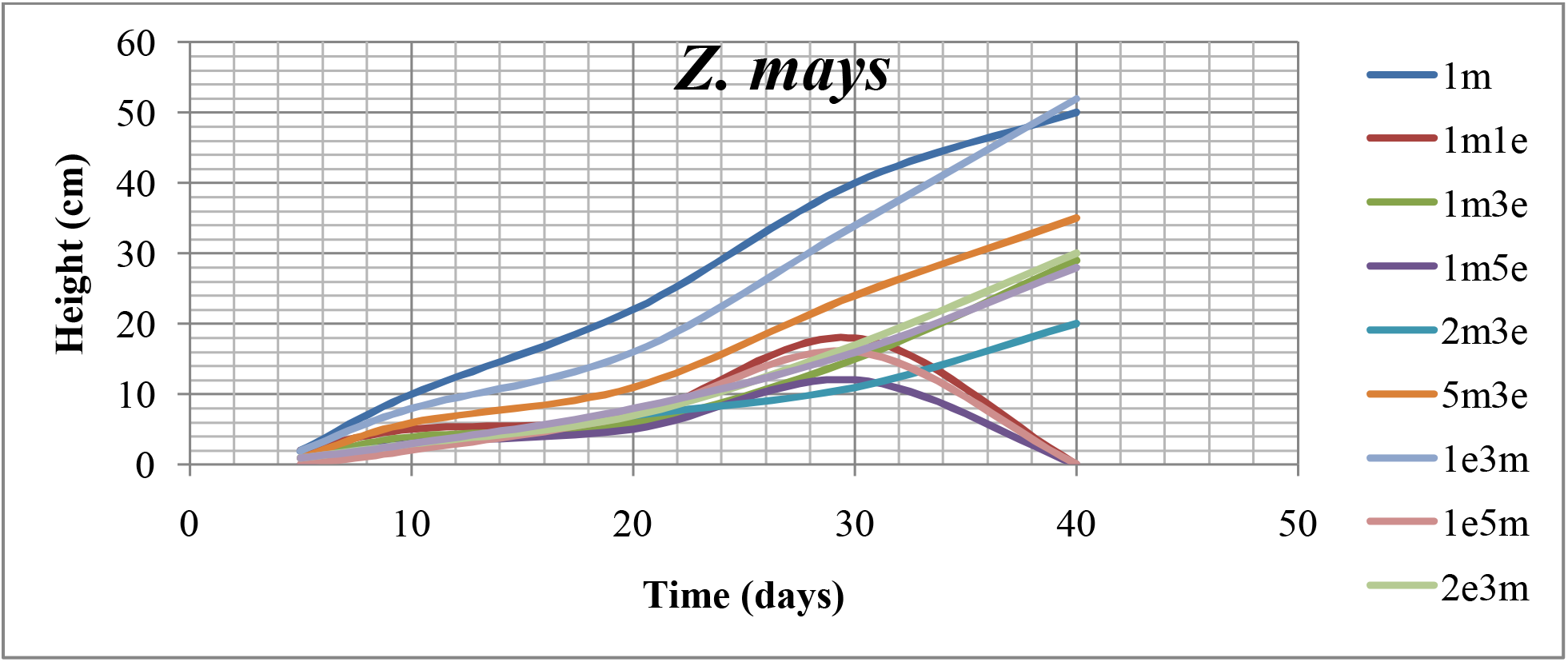
Effects of interaction between *Z. mays* and *E. indica* on plant height of *Z. mays* (The plant interactions are represented on right hand side of graph, indicating number of *Eleusine* plants (e) in interaction with maize (m) plants)

Number of leaves of plants of *E. indica* and maize plants in growth interaction have been presented on Fig. 2(a) and Fig. 2(b) respectively. Results showed significant growth suppression in *E. indica* when the weed was grown in a mix with more maize plants, even though such maize plants did not do better either. This may likely show the competitive strength of the maize plant as well. There were 5 leaves per plant in the weed at the 5^th^ day after growth interaction. At the 40^th^ day however, there were differences in number of leaves in each plant depending on the crop-weed mix ratio; 32 in 1e, 13 in 1e3m, 12 in 2e3m and 20 in 5e3m respectively. Similarly, in the maize (Fig. 2b), increase in number of leaves was inhibited by growth interaction with the weed, especially for those where there were more weeds in interaction than the maize plant. This may likely mean that plant density can affect weed invasion. On day 40, number of leaves was 5 in 1m, 2 in 2m3e, 4 in 2e3m and none in 1m1e, 1m5e, and 1e5m because these plants had died out just after day 30. This aligns with the work of Nwokoclia (1983) who reported that number of maize leaves faced 50% loss due to weed infestation.

**Fig. 2(a):**
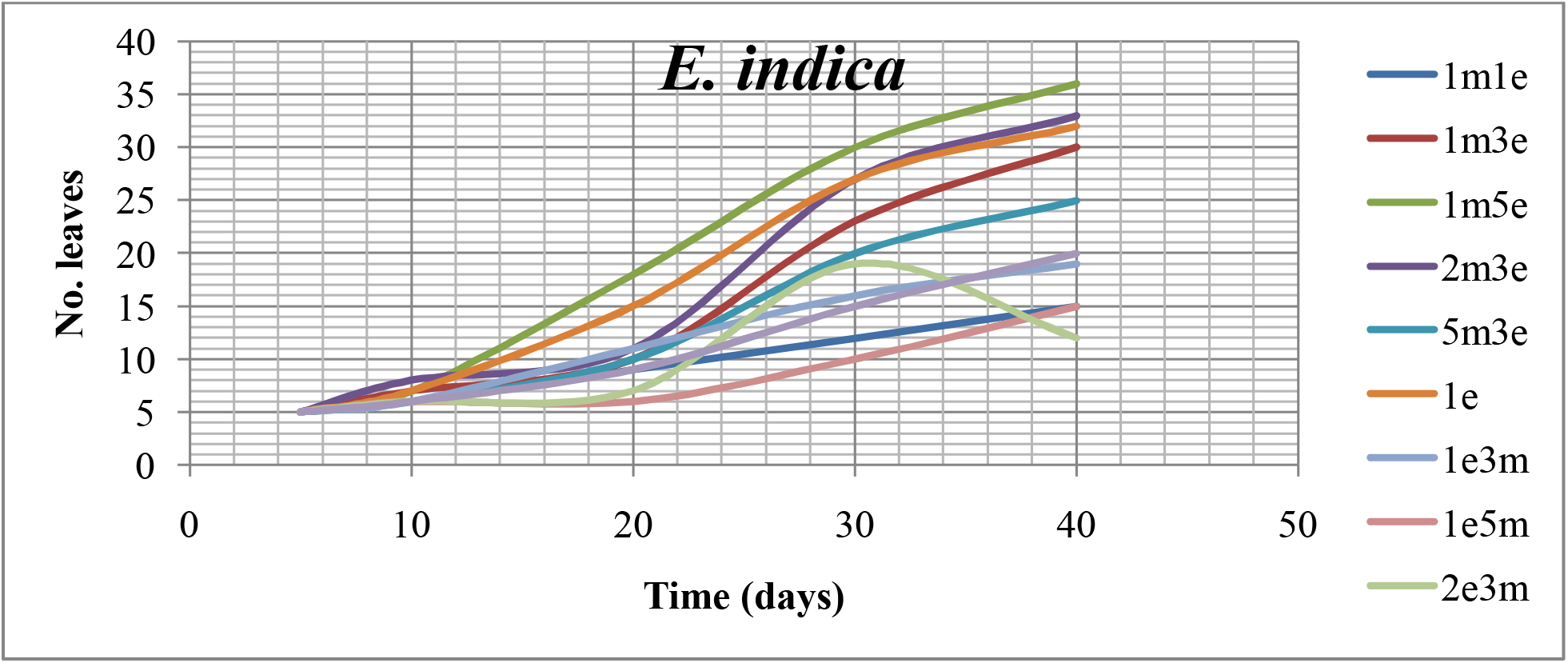
Effects of interaction between *Z. mays* and *E. indica* on the development of number of leaves of *E. indica* (The plant interactions are represented on right hand side of graph, indicating number of *Eleusine* plants (e) in interaction with maize (m) plants)

**Fig. 2(b):**
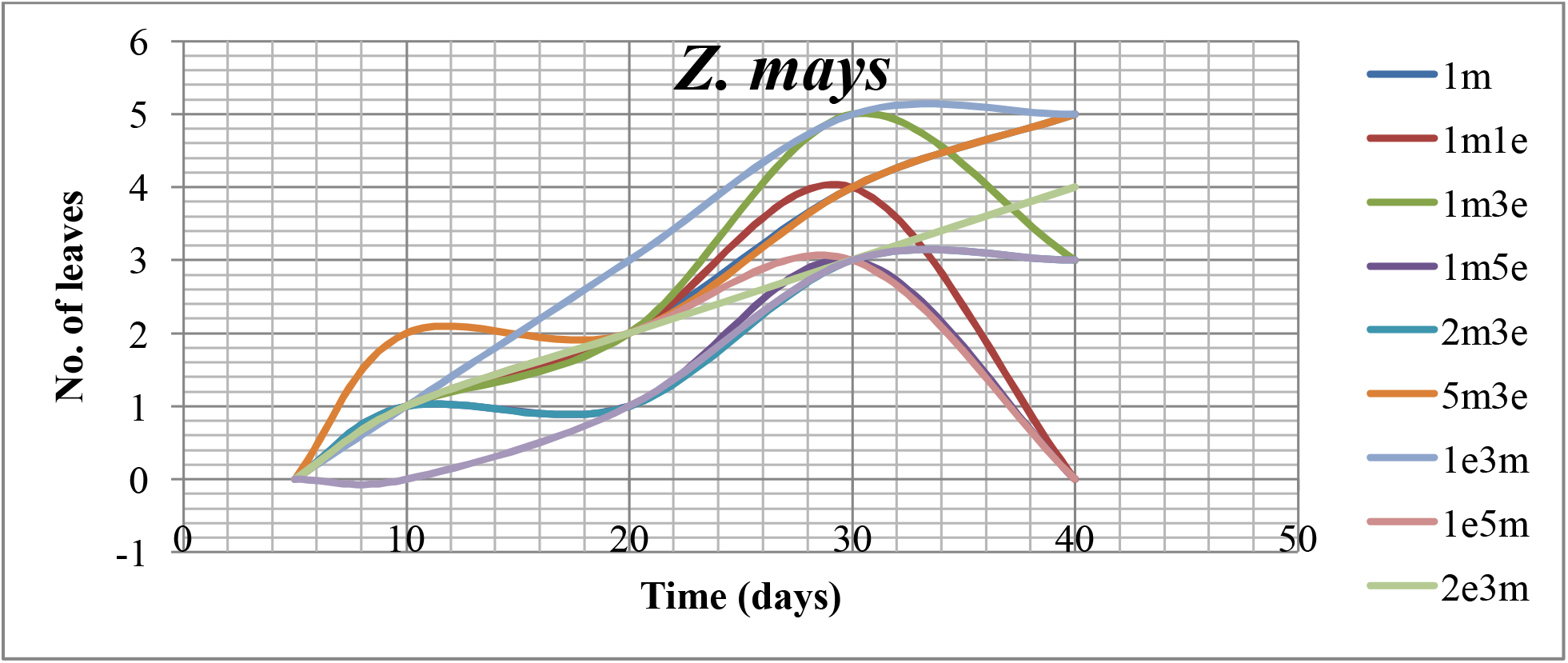
Effects of interaction between *Z. mays* and *E. indica* on number of leaves of *Z. mays* (The plant interactions are represented on right hand side of graph, indicating number of *Eleusine* plants (e) in interaction with maize (m) plants)

For number of tillers, all weeds had just one tiller each as at the 5^th^ day of growth interaction with maize plants (Fig. 3). However, there was increase in number of tillers at the 40^th^ day, depending on the weed-crop mix. The (1m1e) and (5e3m) mix had the highest tillers (8), while the (1e) had the lowest. This showed that as the competition intensifies, both weed and crop develops more tillers. This may be due to increase in soil water level (Bell, 1991).

**Fig. 3:**
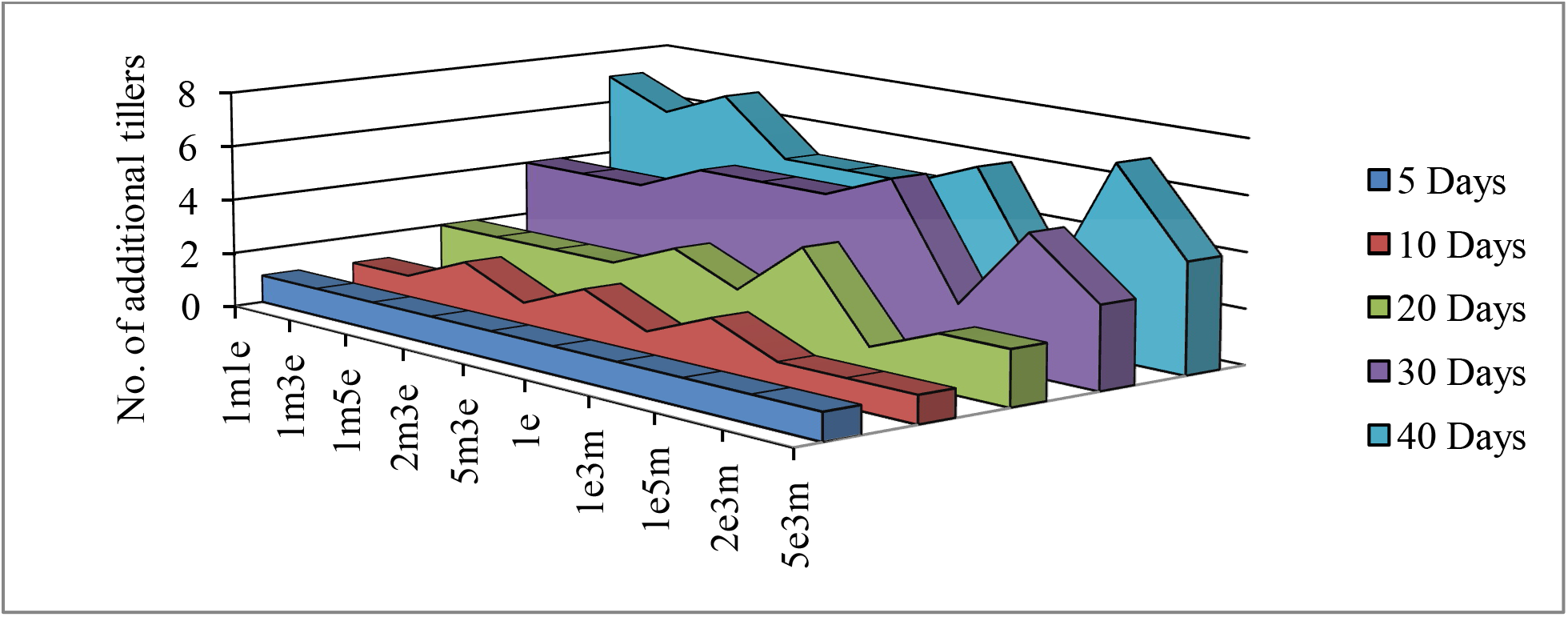
Effects of interaction between *Z. mays* and *E. indica* on gaining of additional tillers by *E. indica* plant(The plant interactions are represented on right hand side of graph, indicating number of *Eleusine* plants (e) in interaction with maize (m) plants)

Figure 4 shows the chlorophyll content index of test plants at 40 days after interaction. For *E. indica* plants, CCI was 1.4cci in 1m1e as against 1.7cci in 1e (Fig, 4a). However, the maximum CCI (4.4cci) was obtained in the 2m3e mix. For maize, however, chlorophyll content index was higher compared to the weed (2.9 – 8.7 cci) for those plants which were still alive at day 40 (Fig. 4b). The highest CCI of 8.7 was obtained in maize plant sown in the 1e3m mix, whereas the least (2.9 cci) was obtained in the 5e3m mix. Maize plants showed higher CCI than *E. indica*, this may likely be as a result of differences in their photosynthetic pathways. *E*.*indica* under maize interaction has a slightly higher CCI than the control. CCI is used to predict the nutritional level of nitrogen in plants. This result did not follow the work of Godoy *et al*., (2008) who stated that increase in beans-weed density reduces the CCI after 40days of interaction.

**Fig. 4(a):**
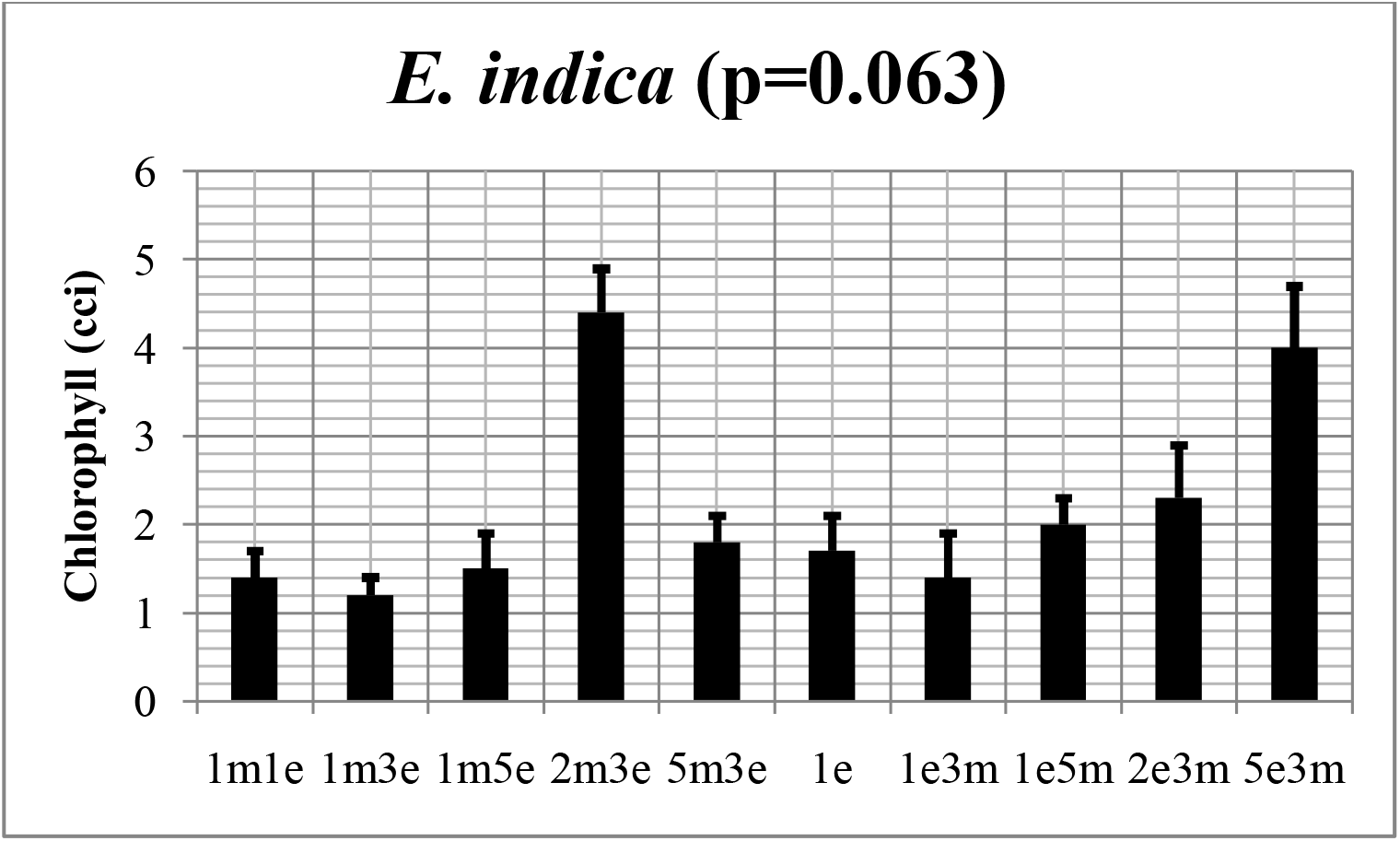
Effects of interaction between *Z. mays* and *E. indica* on foliar chlorophyll of *E. indica* at 40 days after plant interactions(The plant interactions are represented on right hand side of graph, indicating number of *Eleusine* plants (e) in interaction with maize (m) plants)

**Fig. 4(b):**
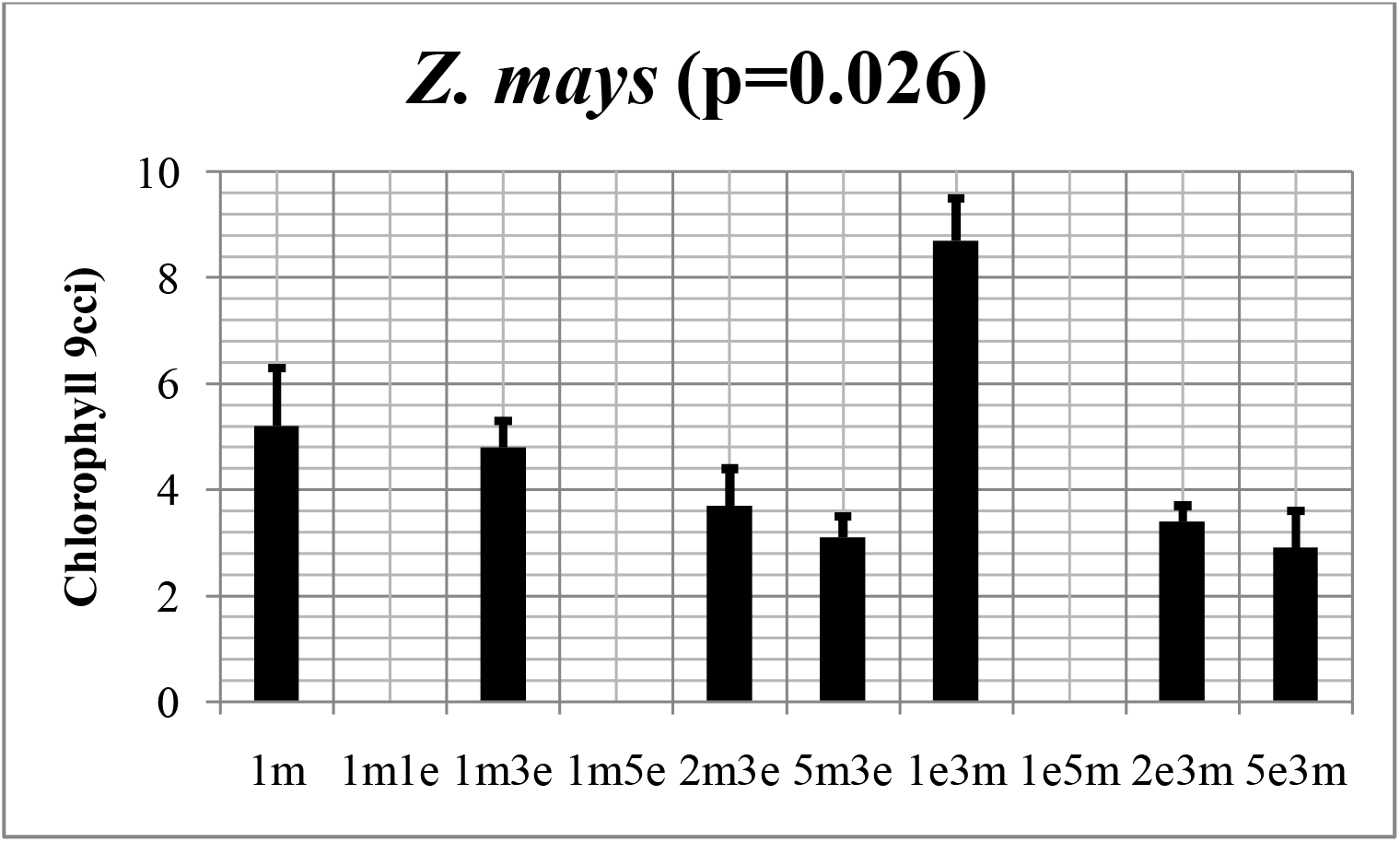
Effects of interaction between *Z. mays* and *E. indica* on foliar chlorophyll of *Z. mays* at 40 days after plant interactions(The plant interactions are represented on right hand side of graph, indicating number of *Eleusine* plants (e) in interaction with maize (m) plants)

Table 1 presented overview results of the experimental parameters assayed in this study. At 40^th^ day after plant interactions, a significant decrease in plant heights of both maize and *E. indica* depending on the ratio of both plants in the crop-weed mix was observed. Similarly, significant decreases in leaf area was reported in *E. indica* (p=0.015) owing to the interaction. In the control, leaf area was 11.4cm^2^, but this decreased significantly to 5.94cm^2^ in the 2e3m mix. Similar decreases in maize leaf area have been reported with leaf area values being as low as 4.32cm^2^ in the 5e3m mix compared to 19.9cm^2^ in the control (1m). The prominent foliar color was yellow green in both plants irrespective of the mix. Chlorosis later set in for both weeds and maize plants. These anti-growth parameters may likely be as a result of the competition for nutrient, water and light between the two mix (Chandramohan*et al*., 2002). Previous work have indicated that competition for environmental resources between maize and weeds cab be attributed mainly to morphological and physiological traits of plants (Didon, 2002). Important trait such as leadf reduction, early vigor, plant height, reduced tillering capacity and initial root and shoot growth are obvious (Betholdsson, 2014).

**Table 1:** Above-ground characteristics of *Z. mays* and *E. indica* upon growth interactions after 40 days

|  | Plant height (cm) |  | Leaf area |  | Presence of foliar chlorosis |  | Foliar colour (prominent) |  | Spike length (cm) |  | Penduncle length (cm) |  |
| --- | --- | --- | --- | --- | --- | --- | --- | --- | --- | --- | --- | --- |
|  | E.I | Z.M | E.I | Z.M | E.I | Z.M | E.I | Z.M | E.I | Z.M | E.I | Z.M |
| 1m | NA | 50a | NA | 19.9a | NA | Yes | NA | green yellow | NA | NA | NA | NA |
| 1m1e | 50b | 0d | 12.2bc | 0d | Yes | No | yellow green | DP | 9.4a | NA | 16.0ab | NA |
| 1m3e | 42bc | 29bc | 6.32bcd | 6.8bc | Yes | Yes | yellow green | green yellow | 5.4ab | NA | 16.3ab | NA |
| 1m5e | 40c | 0d | 14.2a | 0d | Yes | No | forest green | DP | 7.2ab | NA | 12.7bc | NA |
| 2m3e | 69a | 20c | 13.5ab | 11.7b | Yes | Yes | yellow green | green yellow | 6.8ab | NA | 12.4bc | NA |
| 5m3e | 28d | 35b | 4.68d | 11.9b | Yes | Yes | yellow green | green yellow | 7.4a | NA | 17.5ab | NA |
| 1e | 21d | NA | 11.4ab | NA | No | NA | forest green | NA | 2.9b | NA | 11.5bc | NA |
| 1e3m | 29d | 52a | 13.1ab | 21.6a | No | No | forest green | light green | 7.6a | NA | 19.4a | NA |
| 1e5m | 30d | 0d | 19.7a | 0d | Yes | DP | forest green | DP | 6.9ab | NA | 8.5c | NA |
| 2e3m | 23d | 30bc | 5.94cd | 0d | Yes | DP | forest green | DP | 6.5ab | NA | 14.7abc | NA |
| 5e3m | 25d | 28bc | 10.2bcd | 4.32c | Yes | Yes | yellow green | green yellow | 7.2ab | NA | 13.5abc | NA |
| P-value | 0.003 | 0.027 | 0.015 | 0.031 | NA | NA | NA | NA | 0.393 | NA | 0.094 | NA |

NA Not applicable, E.I. *E. indica*, Z.M. *Zea mays*
Means (from 3 replicates) on the same column with similar alphabetic superscripts do not differ from each other ( $p>0.05$ )
The plant interactions are represented on the first column, indicating number of *Eleusine* plants (e) in interaction with maize (m) plants.

Changes in prominent root length were observed in both plants during the interaction (Table 2). Prominent root length was 67.5cm in the control *E. indica* plant (1e) compared to 36.9cm in the 5e3m mix. In the maize plant however, significant reduction in prominent root length was observed from 90.4cm in the control (1m) to 30.5cm in the 2e3m mix. No significant changes in number of primary (7 – 9 roots/plant) and secondary (7 – 12 roots/plant) roots per maize plant was reported (p>0.05). There were generally between 3 and 8 primary roots with secondary branching in both maize and *E. indica*. The presence of possible exudation of organic acids by plant roots during interaction was reported in the present study by Litmus test. Results showed that although there is no evidence of rhizoacidity in the *E. indica* plant, it was however reported in the maize plant only when in the crop-weed mix, which increased intensities when maize plants interacted with more weed in the mix (Table 2). This is consistent with the work of Abdulraheem and Charles (2018), who stated that several allelochemicals are produced in maize, which helps it gain competitive ability and defense against weeds. This may the reason why in some cases, the maize is flourishing better than the weed.

**Table 2:**
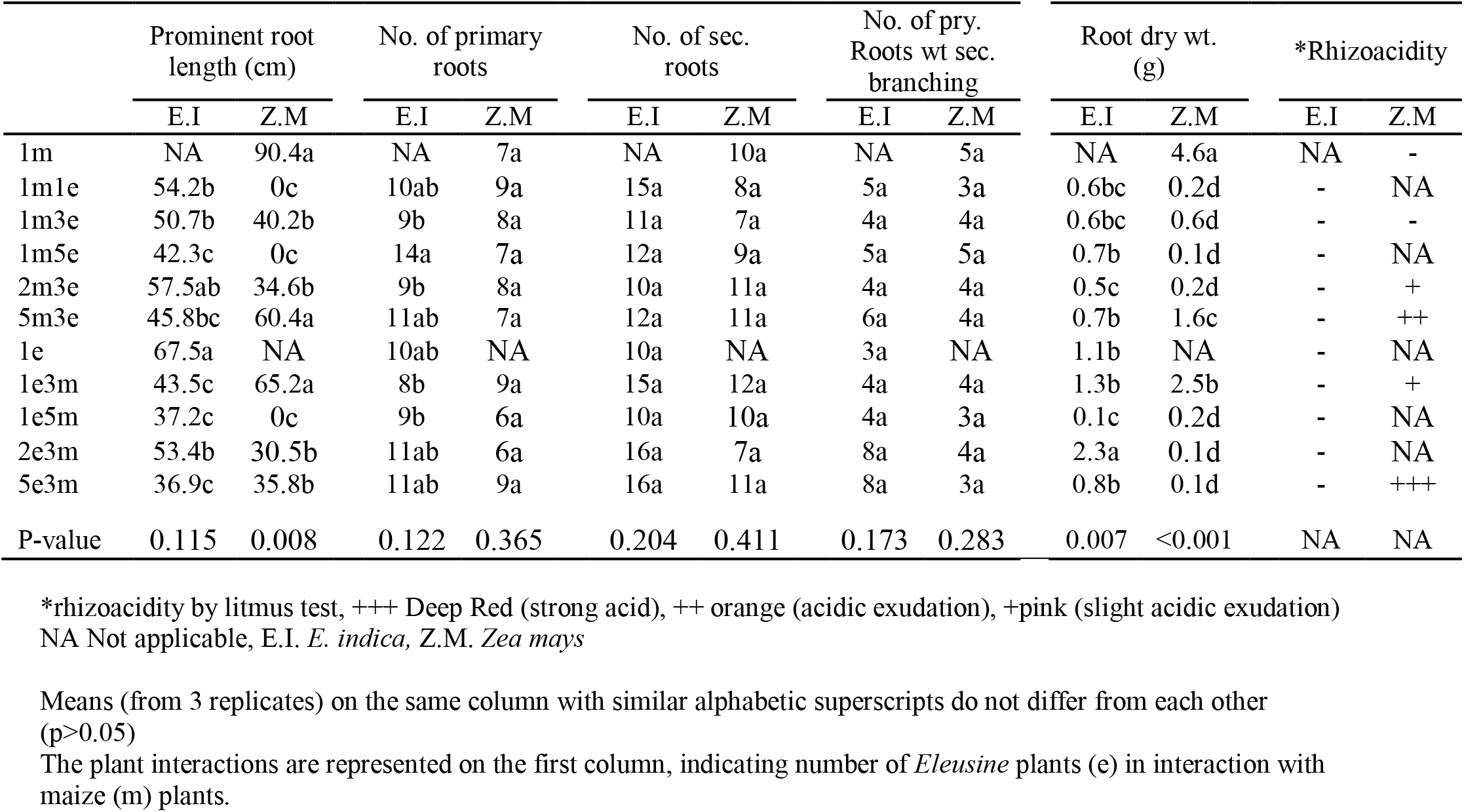
Effects of interaction between *Z. mays* and *E. indica* on root characteristics of both test plants at 40 days after plant interactions

Shoot – root ratios were obtained for both *E. indica* (Fig. 5a) and young maize plants (Fig. 5b). Although the ratio was 1.0 in the control weed (1e), it went as high as 4.5 in 1m1e followed by 3.5 in the 1m3e as well as the 2m3e mix respectively. For maize plant however, the shoot-root ratio was 4.5 in the control (1m), but reduced to as low as 1.5 in the 1e3m mix (Fig. 5b).The results obtained from this study showed that *E. indica* interactions with maize affect maize yield, growth and development. *E. indica* proved to be very harsh and invasive in their competition. However, the maize plant also showed some competitive parameters, even though, some mixtures died before the 40^th^ day of experiment.

**Fig. 5:**
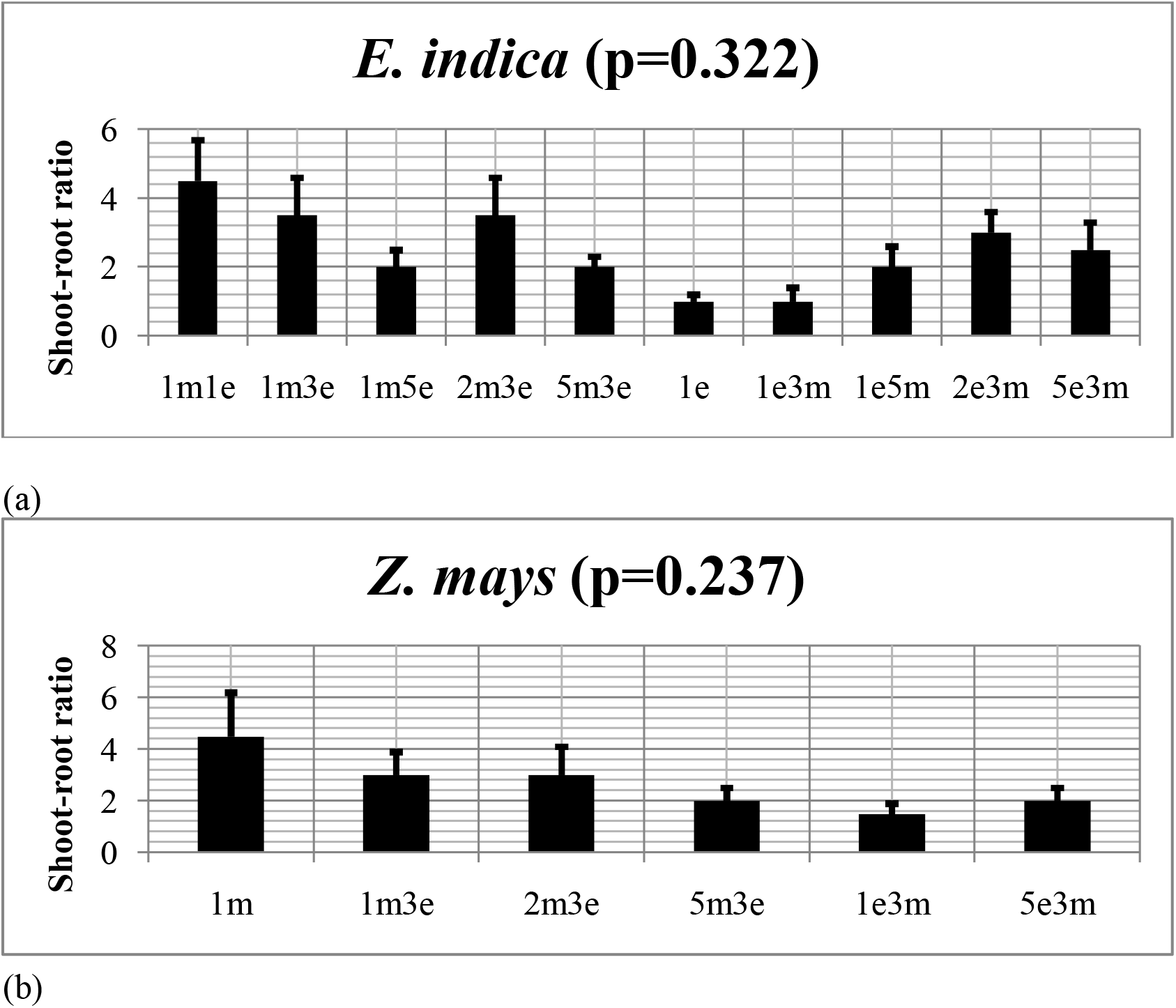
Shoot-root ratio for (a) *E. indica* and (b) *Z. mays* as a result of interaction between both plantsafter 40 days(The plant interactions are represented on x-axis of graph, indicating number of *Eleusine* plants (e) in interaction with maize (m) plants).

## Conclusion

From this study, it can be concluded that a serious competition exist between *E*.*indica* and maize plant when growing in a mixture. Their interaction limit their growth and yield, with more negative effect on the maize plant. Prolong interaction between these two leads to the death of the later. Maize-weed density may also affect this competitive rate. Continuous weeding is necessary for optimum maize yield and food security. Further research should intensify on a more competitive cultivar of maize plant and as well consider the row spacing in comparative study of E. indica-maize competitive strength.

## REFERENCES

Abdulraheem Mukhtar, I. and Charles Eneminyene, F. (2018). Characteristics effects of weed on growth performance and yield of Maize (Zea mays). Biomedical Journal of Scientific and Technical Research. 7(3): 5880-5883. 10.26717/BJSTR.2018.07.001495.

Alpert, P., Bone, E. and Holzapfel, C. (2000). Invasiveness, invisibility and the role of environmental stress in the spread of non-native plants. Perspect Plant EcolEvol Syst. 3:3. 52–66.

Anchal, D., Kapila, S., Anil-Kumar, C., Seema, S., Sanjay, S., Gulsha, M. and Bhagirath, S. (2017). Weed management in rice using crop competitiona review. Crop Protection. 95. 45–52. http://dx.doi.org/10.1016/j.cropro.2016.08.005

Azmi, M. (2000). Journal of Trop. Agric. Food Science. 30:151–161.

Baloch, S., Shah, I. and Hussain, K. (2012). Pakistan Sugar Journal. 17: 13–14.

Bell, A. (1991). Plant Form. Oxford, UK: Oxford University Press. p. 182. 1sbn 0-19-854219-4.

Bertholdsson, N.O. (2014). Variation in allelopathic activity over 100 years of barley selection and breeding. Weed Res. 44: 78–86.

Chandramohan, S., Charudattan, R., Sonoda, R. and Singh, M. (2002). Field evaluation of a fungal pathogen mixture for the control of seven weed grasses. Weed Sci. 50: 204–213.

Companhia Nacional de Abastecimento. (Conab, 2016)Acompanhamento da safrabrasileira de grãos. [accessed on: Mar 2016]. Available on: http://www.conab.gov.br/.

Didon, U.M. (2002). Variation between barley cultivars in early response to weed competition. J. Agron. Crop. Sci. 188: 176–184.

Food and Agriculture Organization (FAO, 2017). Soil and plant testing and analysis. Rome 56/2. 250 p.

Gibson, K.D., Fischer, A.J., Foin, T.C. and Hill, J.E. (2002). Implications of delayed Echinochloa spp. germination and duration of competition for integrated weed management in water-seeded rice. Weed Res.42. 351–358.

Godoy, L.G., Santos, T.S., Boas, R.L. Leite, J.R. (2008). Indice relative de clorofila e o estadonutricionalemnitrogeniodurante o ciclo do cafeeirofertirrigado. RevistaBrasileira de Clencia do Solo. 32:217–227.

Holm, L.G., Pancho, J.V., Herberger, J.P., Plucknett, D.L. (2019). A geographical atlas of world weeds. New York, USA: John Wiley and Sons, 391 pp.

Ikhajiagbe, B., Musa, S. I. and Okeme, J. O. (2019). Effect of changes in soil cation exchange capacity on the reclamation of lead by Eleusine indica (l.) Gaertn. FUDMA Journal of Sciences. 3:4. Pp 176–183.

Kumar, D., and Jhariya, N. A. (2013). Nutriotional, Medicinal and Economical Importance of Corn: A Mini Review. Research Journal of Pharmaceutical Sciences, 2: 7–8.

Nwokoclia, H.N. (1983). Weed interference studies in potato and Maize annual report. National root crops research institute. Pp. 88–93.

Oerke, E.C. and Dehne, H.W. (2004). Safeguarding production-losses in major crops and the role of crop protection. Crop Prot. 23, 275–285.

Ofor, M., Ibeawuchi, O., Oparaeke, M. (2009). Nature and science. 7:45–51.

Ogbeibu, E. A. (2005) Biostatistics: A Practical Approach to Research and Data Handling. Mindex Publishing Co. Ltd., Benin City, Nigerian. pp. 171–173.

Randall, R.P. (2012). A global compendium of weeds. Perth, Australia: Department of Agriculture and Food Western Austalia, 1124 pp.

Shad, R. (2019). Progressive Farming. 7: 10–16.

Shah, J.l., Umed-Ali, L., Ghulam, M.L., Mahmooda, B., Farooque, A.S. (2016). An overview on various weed control practices affecting crop yield. Journal of Chemical, Biological and Physical Sciences. 6:1. 59–59.

Tunku, P. and Ishaya, D. (2012). Effects of cropping pattern and green manure on weed incidence and Productivity of Maize/Soyabean intercrop. Nigerian Journal of Weed Science. 27: 1–9.

Usoroh, N.J. (2003). Field screening of herbicide for weed control in Tomatoes (Lycopersiconesculentum Mill). 13th Annual conference of Weed Science Society of Nigeria. National Cereals Research Institute, Ibadan.

Waterhouse, DF. (2011). Biological control of weeds: Southeast Asian prospects. Canberra, Australia; Australian Centre for International Agricultural Research (ACIAR),v+ 141pp.

Zar, J. H. Biostatistical Analysis. Englewood Cliffs, N. J.: Pretince-Hall, 1974. pp. 471–474.

